# A family of bacterial actin homologues forms a 3-stranded tubular structure

**DOI:** 10.1101/2023.11.07.565980

**Authors:** Julien R.C. Bergeron, Shamar L. M. Lale-Farjat, Hanna M. Lewicka, Chloe Parry, Justin M. Kollman

## Abstract

The cytoskeleton plays a critical role in the organization and movement of cells. In Eukaryotes, actin filaments polymerize into a highly conserved double-stranded linear filamentous structure in the presence of ATP, and disassemble upon ATP hydrolysis. Bacteria also possess actin-like proteins, that drive fundamental cellular function, including cell division, shape maintenance, and DNA segregation. Like eukaryotic actin, bacterial actins assemble upon ATP binding. Longitudinal interactions between bacterial actin protomers along each strand are conserved with eukaryotic actin, but variation in interactions between strands gives rise to striking diversity of filament architectures. Here, we report a family of bacterial actins of unknown function, conserved amongst the *Verrucomicrobiota* phylum, which assembles into a unique tubular structure in the presence of ATP. A cryo-EM structure of the filaments reveals that it consists of three strands, unlike other described bacterial actin structures. This architecture provides new insights into the organization of actin-like filaments, and has implications for understanding the diversity and evolution of the bacterial cytoskeleton.

## Background

Actin is a small, highly conserved protein that plays a crucial role in the structure and function of cells. Actin monomers (G-actin) polymerize to form filaments (F-actin) in the presence of ATP. Actin also possesses ATPase activity, and the filaments disassemble upon ATP hydrolysis and ADP release. This activity is modulated by numerous proteins, which forms the basis for most cellular processes in Eukaryotic cells, including muscle contraction, cell migration, cell division, intracellular transport, and signal transduction^1^. The bacterial cytoskeleton is a complex network of protein filaments that provides structural integrity and shape to the cell, similar to the cytoskeleton in eukaryotic cells. These filaments are often composed of proteins homologous to actin, although in bacteria each protein performs a specific function: MreB form helical filaments that encircle the cell and help maintain the shape of rod-shaped bacteria; FtsA forms a ring-like structure at the cell’s mid-point, known as the Z-ring, and is involved in cell division; ParM transport nascent plasmids to the cell poles; and MamK anchors magnetosome vesicles along a linear track, ensuring their alignment in the cell^2^.

Intriguingly, structural studies have revealed that even though the overall fold and ATPase activity is similar across actin-like proteins, they adopt very distinct filamentous architectures, including parallel, antiparallel, staggered and non-staggered arrangement, which have been proposed to be related to their specific function. Nonetheless, they mainly form two-stranded filaments^3^, although in some case these have been suggested to be part of a higher-order assembly^4-6^.

Here, we report the structure of a previously unreported family of bacterial actin homologues, which forms a rigid, three-stranded, polar filament. This unusual architecture suggests a distinct mechanism of filament assembly, and provides new avenues to our understanding of the evolution of actin-like proteins.

## Results

### A previously uncharacterized family of bacterial actin homologues

Previous studies had demonstrated that each bacterial actin homologue forms a distinct filament architecture^3^. This prompted us to search for previously uncharacterized actin homologues in the bacterial metagenomics database, to identify novel actin-like proteins that could adopt unique filament architectures. We identified a set of proteins, most closely related to MamK^7-10^ (25-30 % sequence identity) (**Figure 1a**) but present in non-magnetotacic bacteria, and therefore presumably with a distinct function. Intriguingly, this protein is highly conserved in the *Verrucomicrobiota* phylum, a poorly characterized family of anaerobic bacteria ubiquitously present in soil samples, as well as in the gut microbiome^11^. Sequence analysis of this protein family confirms that it possesses the hallmarks of actin-like homologues, including conserved residues for ATP binding and hydrolysis (**Figure S1**), and structure prediction confirmed that it likely adopts an actin-like fold. Nonetheless, one distinctive feature of this family is the presence of an additional 30-50 amino-acid extension at the N-terminus (NTD) (**Figure S1**), which is predicted to be unstructured and is not conserved in sequence, but which is not present in other actin-like proteins.

**Figure 1:**
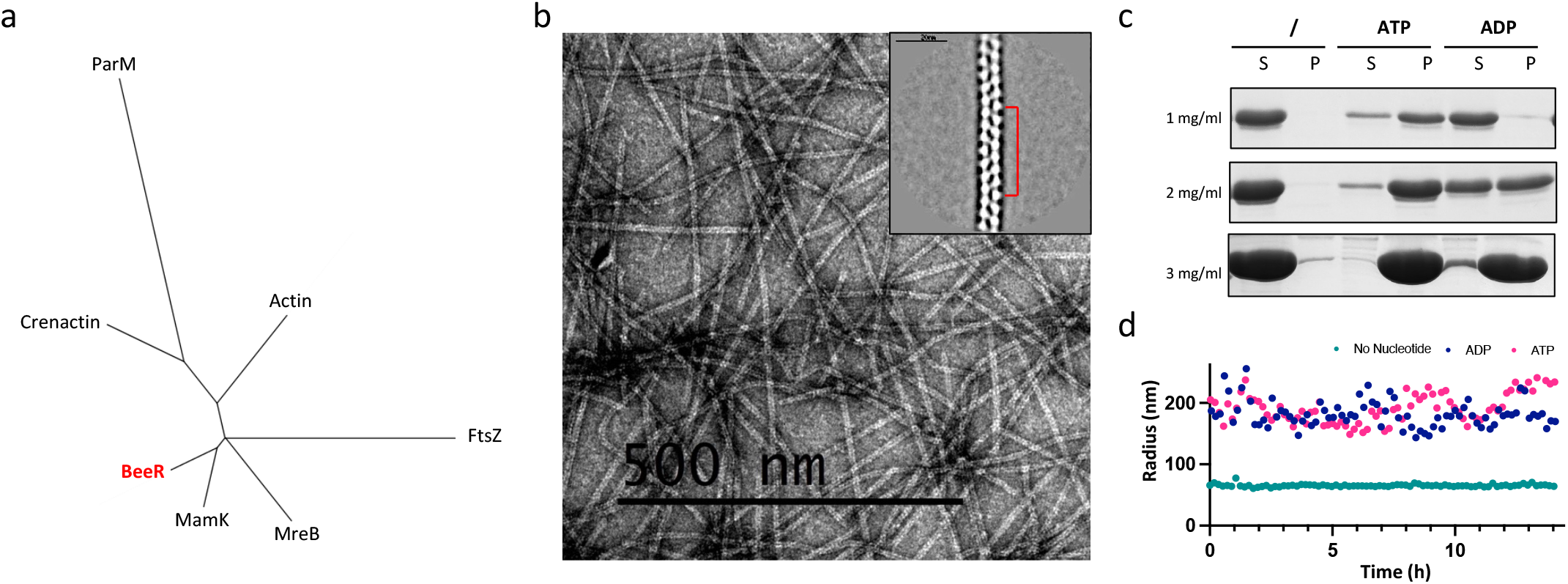
A novel Bacterial actin homologue family forms a three-stranded filament. **a)** Unrooted phylogenic tree of the main families of actin orthologue. BeeR, which is most closely related to MamK, is in red. **b)** Negative-stain TEM micrograph of BeeR filaments in the presence of ATP. Representative 2D classes are shown in insert, revealing a rail tracks-like architecture, with a 7-subunit repeat shown in red. **c)** Pelleting assay for BeeR, at various protein concentration. Monomeric protein is present in the supernatant (S), whereas polymers are found in the Pellet fraction (P). BeeR is primarily monomeric in the absence of nucleotide, and polymerizes in the presence of ATP, as well as ADP at high protein concentration. **d)** Dynamic Light Scattering measurement of BeeR polymerization, in the absence of nucleotide(green), with ATP (Blue), or with ADP (pink). For both nucleotides, no depolymerization is observed over ∼12 h under the conditions employed in this assay.

In order to characterize this newly-identified family, we purified one representative orthologue, from the bacterium *Opitutus terrae*^*12*^ (**Figure S2a**). We then used Negative-stain Electron Microscopy (EM) analysis to verify if this protein formed filaments in the presence of ATP. As shown in **Figure 1b**, indeed this protein forms long, well-ordered filamentous structures, which appeared as twisted rail tracks, very distinct from that of any other actin-like proteins. Notably, these filaments are highly rigid, very long (up to several μm), and 2d classification of these filaments suggested that it adopts a rail tracks-like architecture, consisting of two strands rotating on their individual axis, and with clear 7-subunit repeats (**Figure 1b, insert**). This prompted us to name this family of actin homologues the Bacterial elongated entwined Rail-like protein (BeeR).

A pelleting assay confirmed that BeeR is soluble in isolation, but polymerizes in the presence of nucleotide (**Figure 1c**), demonstrating its actin-like property. We note that at high protein concentration (∼ 3 mg/ml, or 0.1 mM), we observed a similar propensity for polymerization in the presence of both ATP and ADP. However, at lower protein concentration (∼1 mg/ml, or 25 μM), polymerization is vastly more efficient with ATP compared to ADP in this assay (**Figure 1c**). Furthermore, we did not observe significant effects of salt or nucleotide concentration on oligomerization propensity in this assay (**Figures S2b, S2c**). Intriguingly, during the purification process, we also observed that at high concentration, BeeR aggregates at 4°C in the absence of nucleotide (**Figure S2a**), and this aggregation is reversible, with the protein eluting as a monomer in gel filtration at 20°C. We postulate that this indicates a propensity for BeeR to form filaments at low temperature in the absence of nucleotide, however as this aggregation was only observed at high protein concentration, we could not verify whether the aggregates correspond to filaments by negative-stain EM, where required protein concentration is much lower.

In order to characterize the kinetics of filament assembly, we employed Dynamic Light Scattering (DLS) to monitor the oligomerization of this protein over time. As shown on **Figure 1c**, we observed very fast oligomerization, within minutes, with no depolymerization observed within the ∼12h of this experiment. Similar kinetics as observed with both ATP and ADP, however as indicated above, under the conditions of this assay BeeR has an equal propensity to oligomerize with both nucleotide, and therefore we cannot conclude if this corresponds to low nucleotide hydrolysis.

### Cryo-EM structure of the BeeR filament

We next used cryo-EM to gain further insights into the molecular structure of the BeeR filament. As shown in **figure S3a**, BeeR filaments were readily incorporated into ice in cryo-EM grids, which allowed us to collect a cryo-EM dataset of this sample. Following particle picking, we performed 2D classification (**Figure S3b**), which confirmed the rail track-like appearance observed in negative-stain (see above). Based on this, we initially assumed that it corresponded to a two-stranded architecture, but early attempts at determining this structure, using a range of helical symmetry parameters, were unsuccessful.

To identify the helical symmetry of the BeeR filament, we collected tilt series for this sample. In the reconstituted tomograms (**Movie S1**), BeeR filaments appear clearly hollow, indicating that our interpretation of the 2D classes were wrong, and BeeR actually forms a tubular structure. Based on this observation, we were able to obtain an initial 3D reconstruction, using a hollow tube as a starting model. This in turn permitted us to determine the helical symmetry parameters of the BeeR filament, which allowed us to refine this structure by helical reconstruction, to 3.1 Å resolution (**Figure 2b-d, Figure S3c-e and S4, Table S1**). Unexpectedly, this revealed that BeeR adopts a three-stranded, parallel, right-handed, staggered filament architecture (**Figure 2c**). The diameter of this filament, at ∼ 80 Å, is significantly larger than most other actin-like proteins, and it includes a large (∼ 25 Å-wide) cavity at its core (**Figures 2b, movie S2**), responsible for its rail track-like appearance observed in 2D classes.

**Figure 2:**
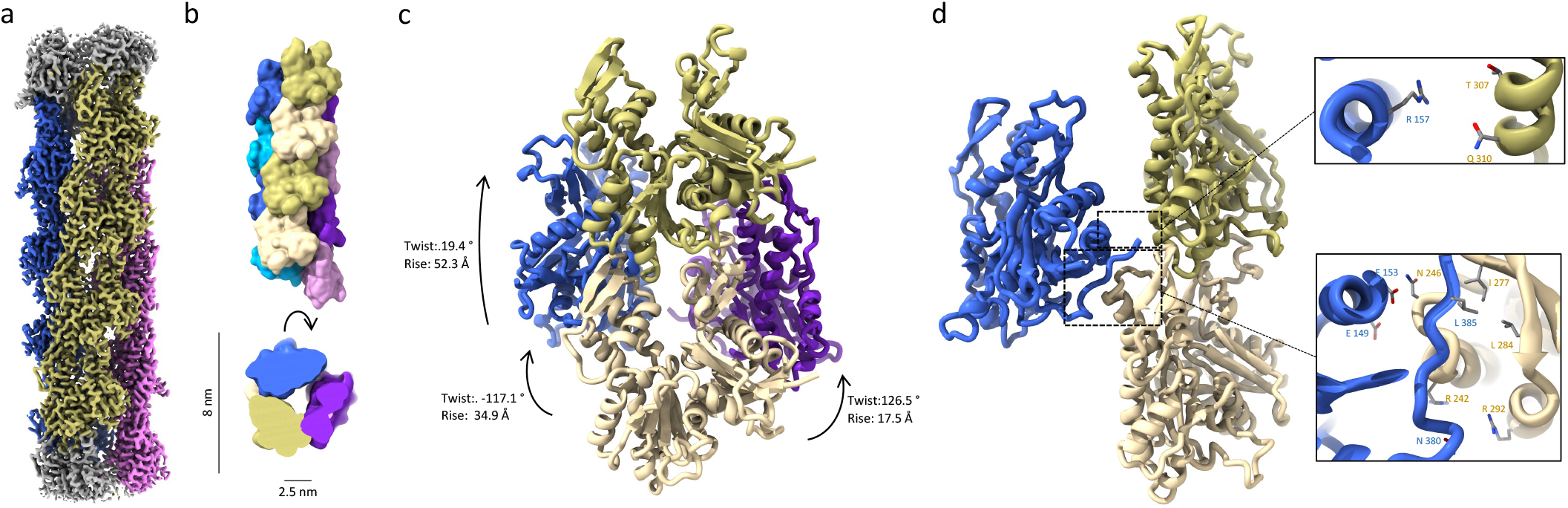
Structure of the BeeR tubular filament. **a)** Cryo-EM structure of the BeeR filament, to 3.1 Å resolution. The three strands are colored in dark blue, khaki, and magenta, respectively. **b)** Surface representation of the BeeR filament atomic model, colored as in **a)**, from the side and top. The diameter of the filament, and internal cavity, are indicated. **c)** Helical symmetry in the BeeR filament. Four adjacent BeeR subunits as shown, illustrating the various helical symmetry parameters that can be used to described this helical structure. **d)** Lateral contact across the strands are shown, with close-up views of the main interacting residues on the right.

The helical arrangement of BeeR can be described in multiple ways: (1) as a 3-start helix, with a rise of one subunit (∼ 52 Å) in a right-handed filament with a twist of ∼ 19 °; (2) as a 2-start helix in a left-handed filament, with a rise of ∼ 35 Å and a twist of ∼ -177 °; or (3) as a 1-start helix in a right-handed filament, with a rise of ∼ 17 Å and a twist of ∼ 126 ° (**Figure 2c**). These are equivalent in terms of subunit arrangement, but we note that representation (3) was used as helical parameters for the cryo-EM map reconstruction, as it has the lowest rise and therefore maximizes the number of individual protomers averaged in our reconstruction.

The structure of the BeeR monomer is consistent with that of an actin-like protein, with the canonical domains Ia, Ib, IIa and IIb adopting a U-shaped arrangement, and the nucleotide buried between domains Ib and IIb^1^ (**Figure S4c**). It is most similar to its closest homologue of known structure, MamK^7,8^, with a RMSD of 1.3 Å for aligned atoms (**Figure 3a**). Within the BeeR filament, the longitudinal interface along strands is largely similar to other actin-like filaments (**Figure 3b**), with domains Ib and IIb making extensive contacts with domains IIa of the subunit above. Notably, its strand arrangement resembles mostly that of MamK, but also actin, both of which have similar interfaces, but with a slightly more twisted filament in the later. The N-terminal domain, which we identified as a hallmark of the BeeR family (see above and **figure S1**) is not resolved in our map, supporting its intrinsically disordered nature.

**Figure 3:**
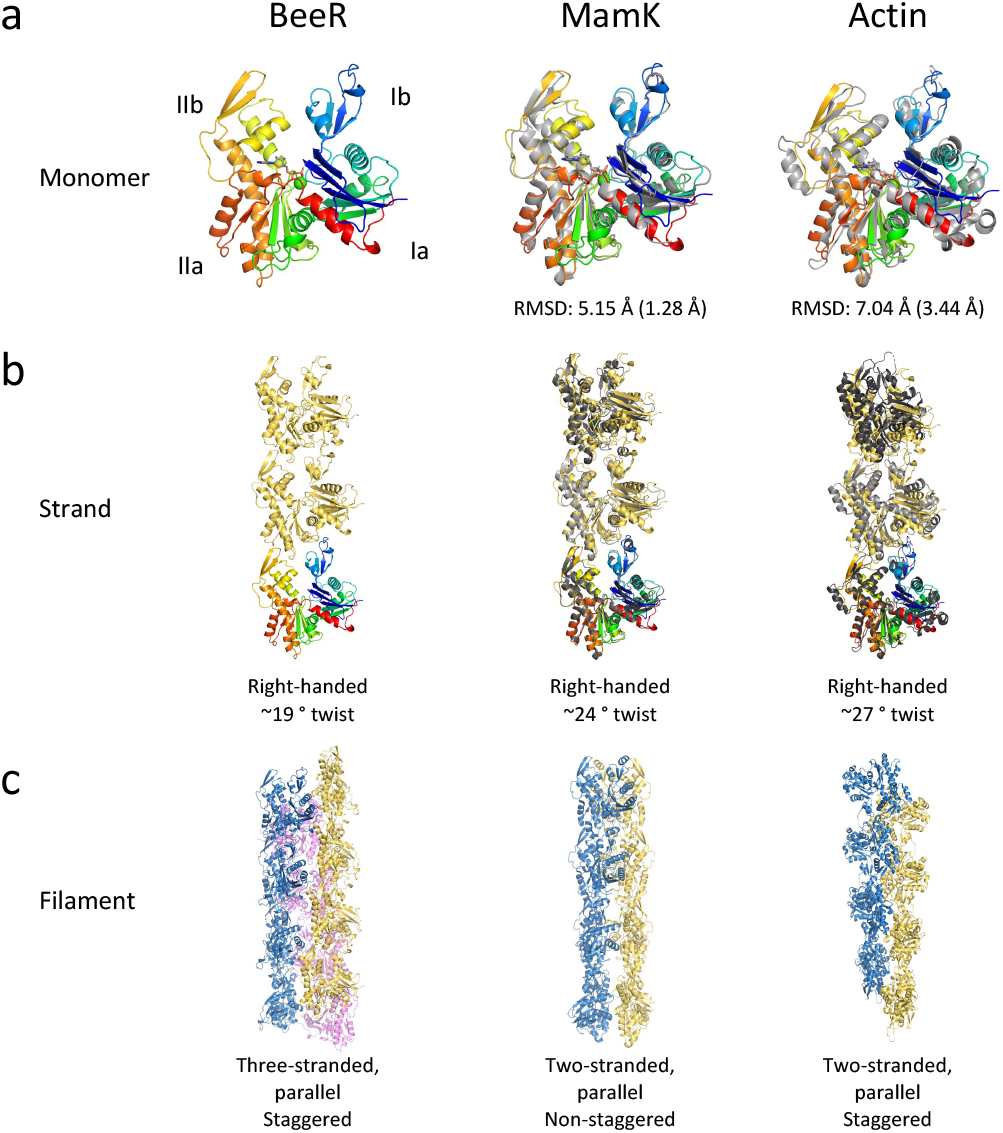
Comparison between BeeR and other bacterial actins. **a)** A monomer of Beer is shown, in rainbow coloring, on its own (left) and overlaid on a MamK monomer (middle) or actin monomer (right) in grey. The RMSD for MamK or Actin is indicated, with the RMSD for CA only in brackets. **b)** Three adjacent BeeR subunits are shown in yellow, on its own and aligned to MamK and actin as is a). The helical parameters for each is indicated. **c)** Full filament structure for BeeR, MamK and actin, colored as in figure 1.

In contrast, the cross-strand contacts in the BeeR filament are strikingly distinct to that of all other actin-like filaments. Each protomer makes contact with two consecutive subunits of the adjacent strand, mainly alongside the 2-start helix interface, as shown on **figure 2h**. Contacts with the top subunit are minimal, with an interface area of ∼35 Å^2^, and consist of a hydrogen bond network between Arg 157 of one subunit, and Thr 307 and Gln 310 in the adjacent subunit. In contrast, the contact with the bottom subunit are extensive, forming a contact surface area of ∼ 450 Å^2,^ and consist of multiple hydrogen bonds, between Glu 149/Glu 153 and Asn 246, and between Arg 242/Arg 252 and Asn 380. In addition, a hydrophobic residue at the C-terminus, Leu 385, is buried in a hydrophobic pocket of the adjacent subunit consisting of Ile 277 and Leu 284.

### The N-terminal tail of BeeR prevents bundling and enhances filament stability

As mentioned above, a hallmark of the BeeR family is the presence of a predicted disordered NTD. We therefore sought to assess if this NTD was involved in filament formation. As shown in **Figure 4a**, a construct lacking the NTD (BeeR_ΔN_) still retained its capacity to polymerize, and the resulting filaments adopt the same rail track-like morphology (**Figure 4b)**. However, we observed that those filaments have a very high propensity to form larger bundles, consisting of multiple filaments sticking together (**Figure 4a**). These were not observed with the wild-type BeeR protein. Based on this, we propose that the NTD of BeeR forms a “fuzzy coat” (**Figure 4c**) acting as a repellant, and preventing the formation of bundles that might alter their cellular function.

**Figure 4:**
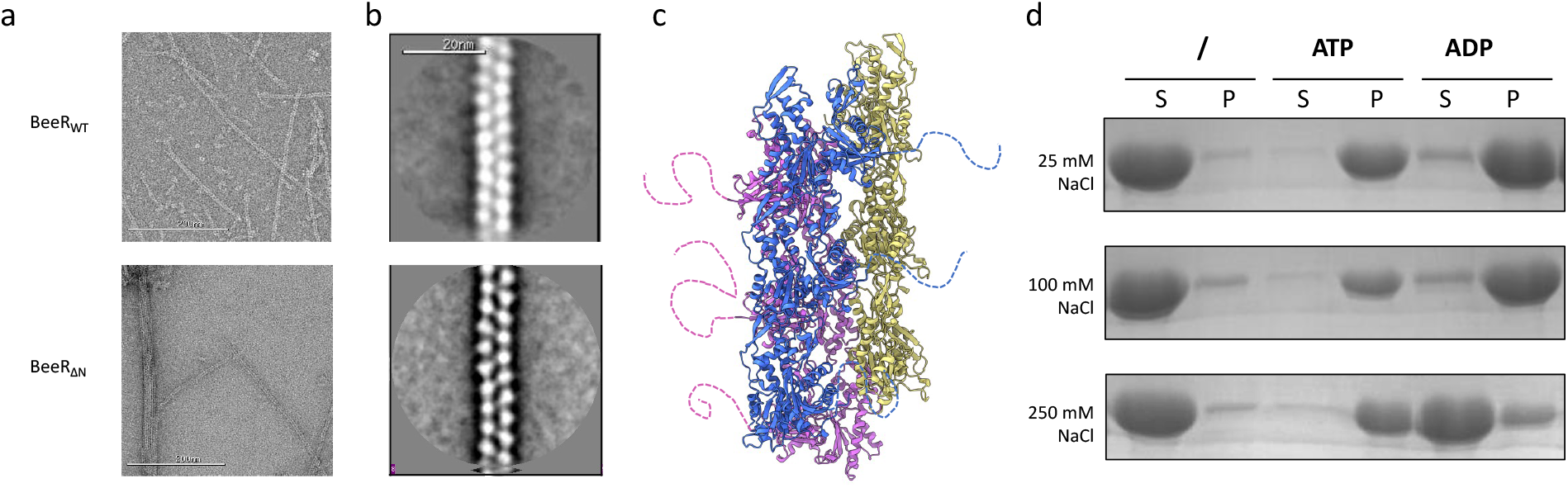
The BeeR filaments bundle in the absence of the N-terminal unstructured domain. **a)** Negative-stain TEM micrograph of BeeR filaments obtained with the WT protein (top) and with a construct lacking the N-terminal domain (bottom). **b)** 2D classes of filaments from the corresponding proteins. **c)** Atomic model of the BeeR tubular filament, with the N-terminal tail indicated in doted lines, forming a fuzzy coat that prevents bundling. **d)** Pelleting assay for BeeR_ΔN_, at different salt concentrations. At high salt concentration, ATP but not ADP promotes BeeR_ΔN_, confirming a role for charge of the N-terminal tail in oligomerization.

Previous studies other bacterial actin-like homologues, including MamK^9^ and AlfA^13^, revealed a strong propensity for bundling, that alters their ATPase activity, and which is modulated by ionic strength. We therefore sought to verify if the propensity to polymerize was altered by the N-terminal tail. To verify this, we employed the pelleting assay described above. As shown on **figure 4d**, at low ionic strength, BeeR_ΔN_ oligomerizes with both ATP and ADP, similar to BeeR. However, at physiological ionic strength, BeeR_ΔN_ only oligomerizes with ATP, and is predominantly monomeric with ADP. This demonstrates that the bundling induced by the deletion of the N-terminal domain destabilizes the oligomerization propensity, in a salt-dependent fashion.

## Discussion

The function of BeeR is not known, and its expression or function has not been investigated to date. Nonetheless, the fact that it is conserved across the phylum suggests that it likely plays a role in *Verrucomicrobiota* biology.

Both the architecture of the BeeR filaments, and its biochemical properties, hints at a function distinct to that of other bacterial actin homologues. The diameter of its cavity (∼ 2.5 nm) makes it unlikely that it is involved in solute or macromolecular transport within the tube. Instead, we hypothesize that the three-stranded tubular structure formed by BeeR provides a much more rigid filament than that of other actin homologues, as supported by 2D classes (**Figure 1b, S3b**). This likely will have an impact on its capacity to perform its function within the cell.

It is noteworthy that most of the residues involved in the tubular architecture of BeeR are conserved among orthologues, but not in MamK or MreB (**Figure S1**). This suggests that the three-stranded filament architecture is a feature of this protein family, and not unique to the particular *O. terrae* orthologue characterized here. Most notably, the majority of the cross-strand interaction occurs between helix a3 and the C-terminal ∼ 10 residues. In contrast, we note that the loop forming the cross-strand interface in MamK (residues 80-85) is not present in any of the BeeR orthologues (**Figure S1**), confirming that they cannot adopt a similar 2-stranded architecture.

The structure of the BeeR filament provides novel insights into the molecular diversity of actin-like proteins, and adds to the diverse structural architectures that can be formed by this family. It uncovers novel paths towards our understanding of the evolution of the actin fold.

## Materials and Methods

### Protein expression and purification

The *DNA* sequence for the *O. terrae* BeeR orthologue, codon-optimised for expression in *E. coli*, was cloned into pET21a. The protein was over-expressed in BL21a cells for 5h at 37 °C. Cells were harvested at 6,000 g for 12 min, and resuspended in BeeR lysis buffer (10 mM HEPES pH 7.0, 25 mM NaCl, EDTA-free protease inhibitor tablets). They were then lysed by sonication for 5 mn, debris were spun down at 45,000 g for 20 min. BeeR was purified from the supernatant by two consecutive steps of ammonium sulphate precipitation at 25% saturation, for 1h each. A following gel filtration step was added for analytical purposes, using a Superdex200 column (Cytiva), in BeeR lysis buffer.

The construct for BeeR_ΔN_ was engineered by removing residues 2-34 by site-directed mutagenesis. The protein was purified as described above.

### Oligomerization assays

For pelleting assays, BeeR or BeeR_ΔN_ was diluted to 3mg/ml (unless specified), in BeeR lysis buffer (with additional NaCl added, when specified in the results section), and both nucleotide and MgCl2 were added to 5 mM (unless specified); only MgCl2 was included in the no-nucleotide controls.

For the DLS assay, protein samples were diluted to 3mg/ml in BeeR lysis buffer supplemented with 5 mM MgCl2, and 100 μl was added to three wells in a 96-well microplate (Greiner). Nucleotide at 5 mM final concentration, or water, was added immediately prior to starting the experimend. DLS measurements were performed in a DynaPro III plate reader (Wyatt), every 10 min for 16 hours, at 20 °C.

For negative-stain TEM experiments, samples were diluted to 0.1 mg/ml, incubated with ATP and MgCl2 at 1 mM, and applied to glow-discharged carbon-coated cupper grids. Samples were imaged on a Tecnai G2 spirit TEM microscope (Thermo Fisher), equipped with a Ultrascan 4000 CCD (Gatan). For data processing, ∼ 20 micrographs were collected manually, at a range of defocus values (−0.8 to -2 μm). CTF estimation, particle picking, and 2D classification was performed in Relion 3^14^.

### Cryo-Electron Microscopy

BeeR was diluted to ∼2 mg/ml, and ATP and MgCl2 were added to a final concentration of 1 mM, and applied to holey carbon grids. For cryo-ET, 10nm colloidal gold was also added as fiducial markers. Grids were then plunge-frozen in liquid ethane using a Leica EM GP2 vitrobot (Thermo Fisher).

For cryo-electron tomography, grids were imaged in a Tecnai F20 TEM (Thermo Fisher), equipped with a K2 summit camera (Gatan), using Leginon^15^. 10 tilt series were collected, consisting of 40 micrographs with tilt ranging from -60 to +60, with a 3 ° tilt between frames, and with a total dose of 100 e^-^/Å^2^. For single-particle analysis, grids were imaged in a Glacios TEM (Thermo Fisher) equipped with Flacon IV camera. A dataset of ∼ 4,000 micrographs was collected with EPU. Cryo-EM data was processed with CryoSPARC^16^, and the structure was determined by helical refinement, using a rise of 17.4 Å and a twist of 126.5°.

An initial atomic model of BeeR was generated with the AlphaFold2 Google Collab server^17^, and 36 copies were placed in the corresponding density of the EM map. The obtained model was subject to real-space refinement in Phenix^18^.

All structural figures were made using ChimeraX^19^.

## Acknowledgements

We are grateful to Ertan Ozyamak and Arash Komeili for initially identifying BeeR as a MamK homologue. JRCB acknowledges members of the Bergeron lab for occasionally tolerating their PI in the lab. Cryo-EM data was collected at the Imperial College Electron Microscopy Centre (funded by BBSRC equipment grant BB/V019732/1), and we acknowledge Paul Simpson for support. DLS experiments were performed in the Biomolecular Interactions facility at King’s College London (Funded by BBSRC equipment grant BB/V01966X/1), and we acknowledge Tam Bui for support. This work was funded by grants from the BBSRC (BB/R009759/2) and HFSP (RGY0080/2021) to JRCB, and the National Institutes of Health (R35149542) to JMK.

## Data availability

The electron potential map of the BeeR filament is available in the EMDB, with accession number EMD-18852. The atomic model is available in the PDB, with accession number 8R2N.

## Author contributions

JRCB, SLML-F, HML, CP performed experiments; JRCB wrote the manuscript; JRCB and JMK conceptualized the project.

## Competing interest statement

The use of BeeR and its derivatives for targeted drug delivery is protected by patent PCT/GB2024/052768. This patent is licensed exclusively to Prosemble group LTD, for which JRCB is a co-founder.

## Notes

### Competing Interest Statement

The authors have declared no competing interest.

### Summary of Updates

Additional peletting and DLS assays have been included to expand on the oligomerization property of BeeR (Figures 1c, d, Figure S1). The Cryo-EM section has been expanded to include additional experimental details (Movie S1, Materials and methods). Additional characterization of the BeeRDN mutant has been included (Figure 4c, d).

