## Supplementary figures and tables for "A family of bacterial actin homologues forms a 3-stranded tubular structure"

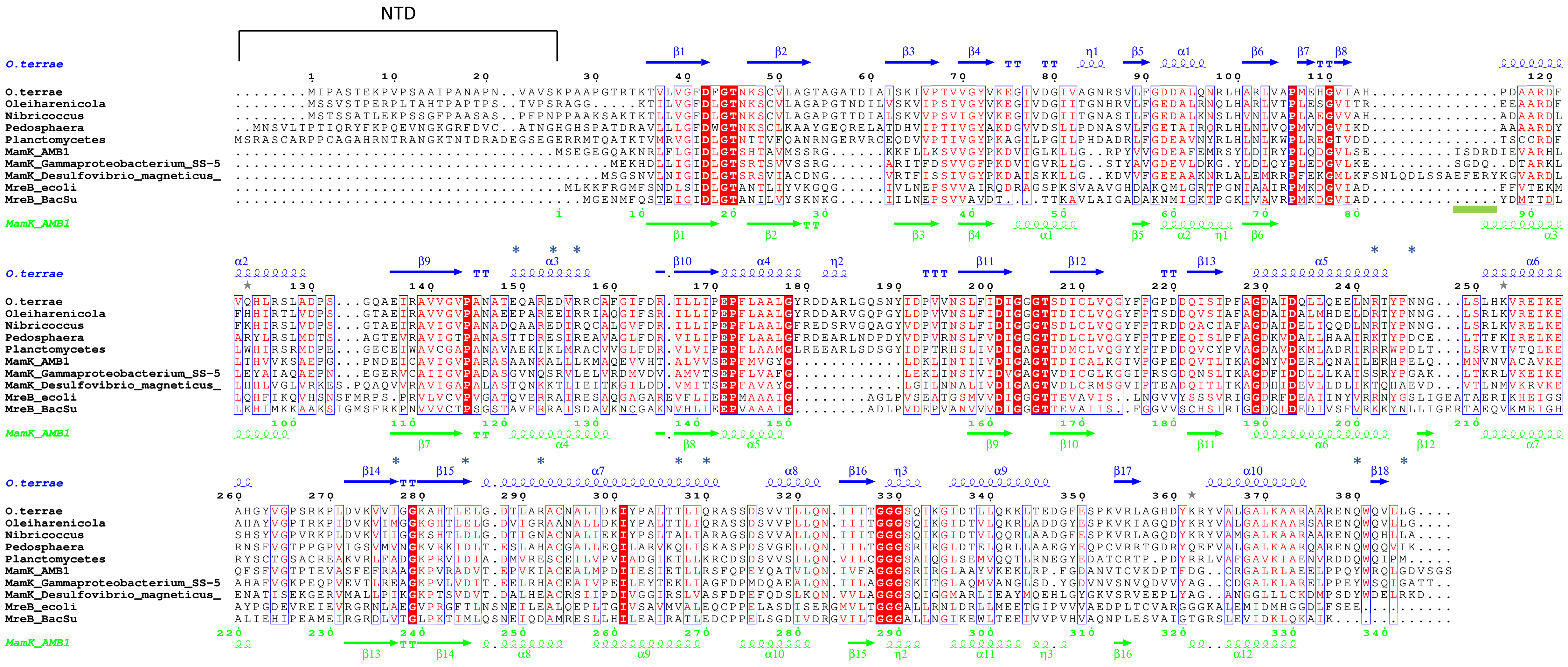


**Figure S1: Multiple Sequence Alignment of selected BeeR orthologues.** BeeR sequences from various bacterial strains, as well as the sequences for several MamK and MreB orthologues, are indicated. Conserved residues are in red boxes, similar residues are in red character. The secondary sequence of BeeR and MamK are shown on top (blue) and bottom (green), respectively. The blue * symbol indicates residues located at the cross-strand interface; the green line indicates the loop forming the cross-strand interface in MamK.

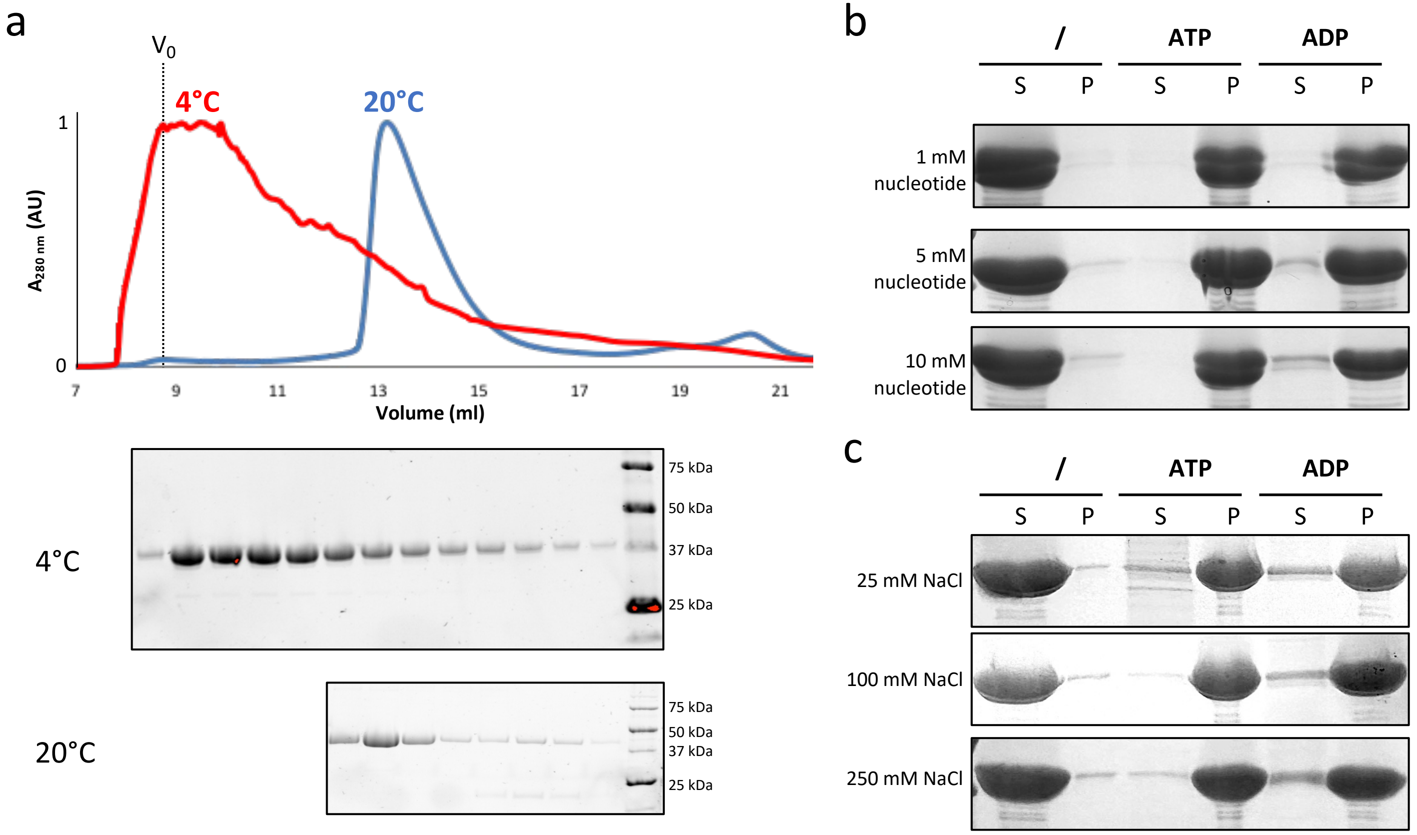


**Figure S1: Oligomerization propensity of BeeR. a)** UV traces for size exclusion chromatography of BeeR, at 4 °C (red) or 20 °C (blue). The corresponding SDS-PAGE gels are shown below. At high concentration, BeeR aggregates at 4 °C, but elutes in a monomeric state at 20 °C. **b)** Pelleting assays for BeeR at various nucleotide concentration, and **c)** pelleting assays for BeeR at various salt concentration. Within the conditions of this assay, nucleotide or salt concentrations do not affect propensity for polymerization.

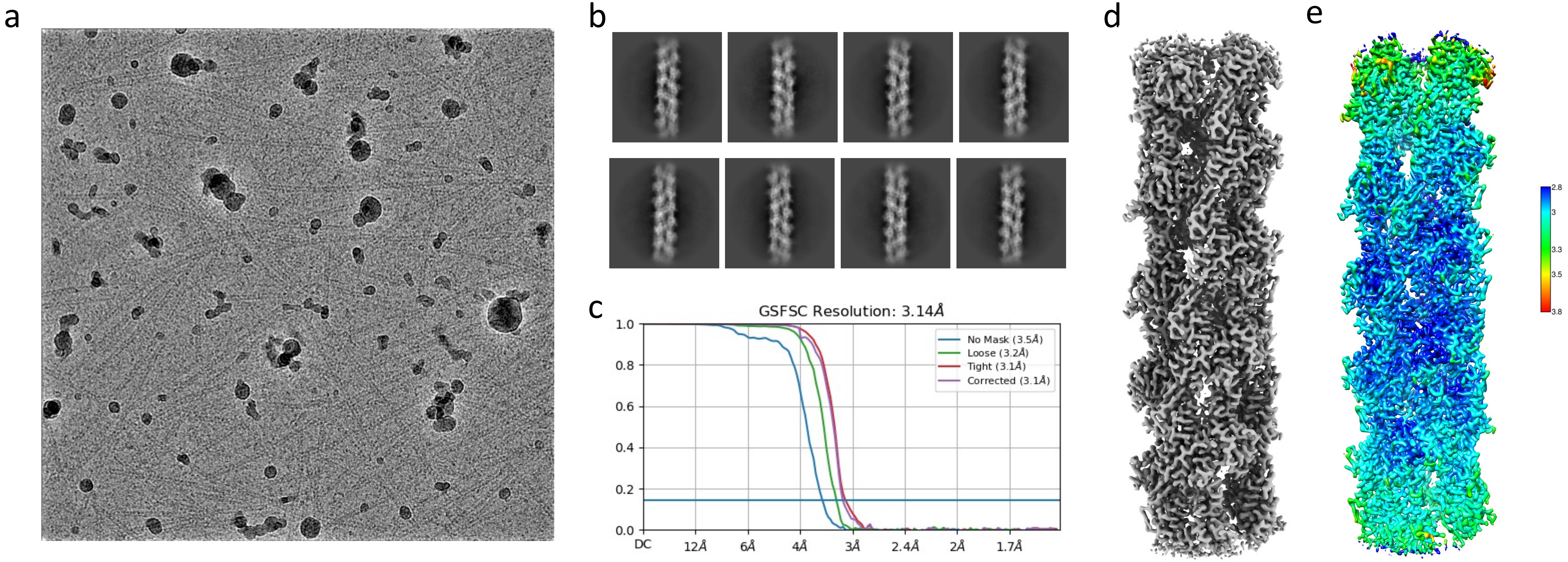


**Figure S3: Cryo-EM analysis of the BeeR filaments. a)** representative electron micrograph, **b)** selected 2D classes, **c)** FSC plot for the 3D refinement, **d)** final map, and **e)** local resolution estimation, showing that the map is mostly in the 2.8 Å – 3.3 Å range.

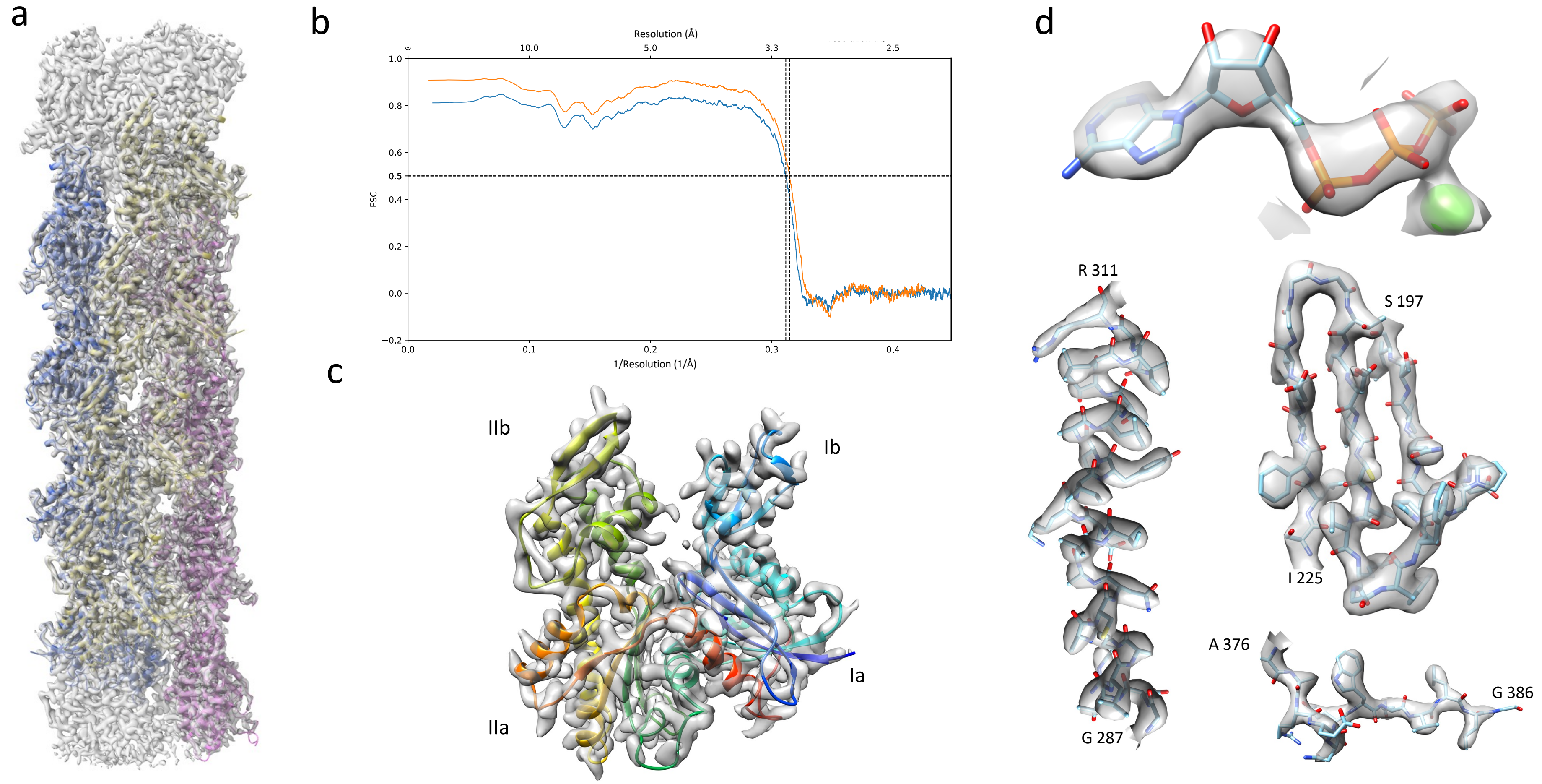


**Figure S4: Atomic model of the BeeR filament. a)** Complete atomic model, colored as in Figure 2, fitted in the electron potential map in grey. **b)** Map-to-model FSC plot for the complete atomic model. **c)** Close-up view of a BeeR monomer, in Rainbow coloring, in the EM map density. The four typical actin domains are indicated. **d)** Representative density for various elements of the structure, including the nucleotide (top), a helix (left), 3-sheet strand (right), and C-terminal loop (bottom), illustrating that the map density matches the reported resolution.

| | | |
|---|---|---|
| **Data Collection** | Microscope | Glacios |
| | Camera | Falcon IV |
| | Voltage | 200 kV |
| | Pixel size | 1.5 |
| | Defocus range (nm) | -0.5 to -2 |
| | Dose | 40 |
| | Micrographs | 4002 |
| **Reconstruction** | Software | CryoSparc |
| | Rise | 17.5 |
| | Twist | 126.5 |
| | Initial particle n. | 2,941,546 |
| | Final particle n. | 882,189 |
| | Resolution | 3.14 |
| **Refinement** | Atoms | 33252 |
| | Protein AA | 4236 |
| | ATP | 12 |
| | Mg | 12 |
| | Map CC | 0.89 |
| | B-factor protein | 29.98 |
| | B-factor ligands | 26.74 |
| | Clashscore | 10.17 |
| | Rotamer outliers (%) | 3.25 |
| | Ramachandran favored | 95.99 |
| | allowed | 4.01 |
| | outliers | 0 |
| | Bond length (A) | 0.004 |
| | Bond angle (deg) | 0.65 |

Table S1: Cryo-EM data collection, processing, and model refinement statistics.
